# How many colours can you see? Real environmental lighting increases discriminability of surface colours

**DOI:** 10.1101/2024.04.23.590719

**Authors:** Takuma Morimoto, João M. M. Linhares, Sérgio M. C. Nascimento, Hannah E. Smithson

## Abstract

Color supports object identification. However, two objects that differ in color under one light can appear indiscriminable under a second light. This phenomenon, known as *illuminant metamerism*, underlies the difficulty faced by consumers of selecting matching fabric or paint colors in a store only to find that they appear not to match under home lighting. The frequency of illuminant metamerism has been evaluated only under single, uniform illuminants. However, in real world conditions, the spectral content of light falling on an object varies with direction (Morimoto et al. 2019), meaning that a surface will sample different spectra depending on its angle within the environment. Here we used computer-graphics techniques to simulate a pair of planar surfaces placed under newly measured hyperspectral illumination maps that quantify the directional variability of real-world lighting environments. We counted the instances of illuminant metamerism that can be solved simply by viewing surfaces tilted to a different direction. Results show that most instances of illuminant metamerism can in theory be resolved for both trichromatic and dichromatic observers. Color deficient observers benefit more than trichromats implying that the directional variability allows the recovery of the ‘missing’ dimension in their colour vision systems. This study adds a new perspective to the classic trichromatic theory of human vision and emphasizes the importance of carefully considering the environments in which biological vision operates in daily life. It is striking that the physical directional variability available in natural lighting environments substantially mitigates the biological limitations of trichromacy or dichromacy.

## Introduction

Human perception of surface color is mediated by light reflected from a surface entering the eye and eliciting signals from retinal cones. Importantly, for a single cone, information about the wavelengths of incoming photons is lost in the resultant neural signal (the so-called principle of univariance (Rushton, 1972)). This is a fundamental limitation imposed at the very first stage of visual processing, which cannot be undone by any higher-level process. Thus, the sensory input to trichromatic human colour vision is limited to triplets of signals from the three classes of cone with different spectral sensitivities (Young, 1802). A monochromatic light at 580 nm elicits a yellow percept, but it is possible to generate a colour that appears identical by balancing the amounts of two monochromatic lights at, say, 520 nm and 620 nm. In a similar way, the lights reflected from two surfaces that have different spectral reflectances can elicit the same triplet of cone signals under one illumination but a different triplet under a second illumination. Past studies have shown that this *illuminant metamerism* occurs relatively frequently, which imposes significant constraints on human color vision (Logvinenko et al., 2014). For dichromats, who possess only two classes of cones, illuminant metamerism can be an even more severe problem (Logvinenko et al., 2015).

In past studies, however, the prevalence of metamerism has been evaluated under an oversimplified environment where surfaces are uniformly illuminated by a single-spectrum illuminant. Recently collected hyperspectral environmental illumination maps (Morimoto et al., 2019) have revealed that natural environments hold significant directional spectral variation; a surface placed in the natural environment samples different illuminant spectra depending on its tilt. This means that a pair of two flat surfaces that appear to have identical surface color at one tilt (Figure 1 A) may be disambiguated simply by adjusting the tilt (Figure 1 B). Here we used computer-graphics techniques to evaluate the frequency of metamerism for planar surfaces placed in a three-dimensional environment defined by the measured illumination maps. Furthermore, we evaluated the extent to which natural directional variability increases chromatic discriminability of surfaces.

**Figure 1.**
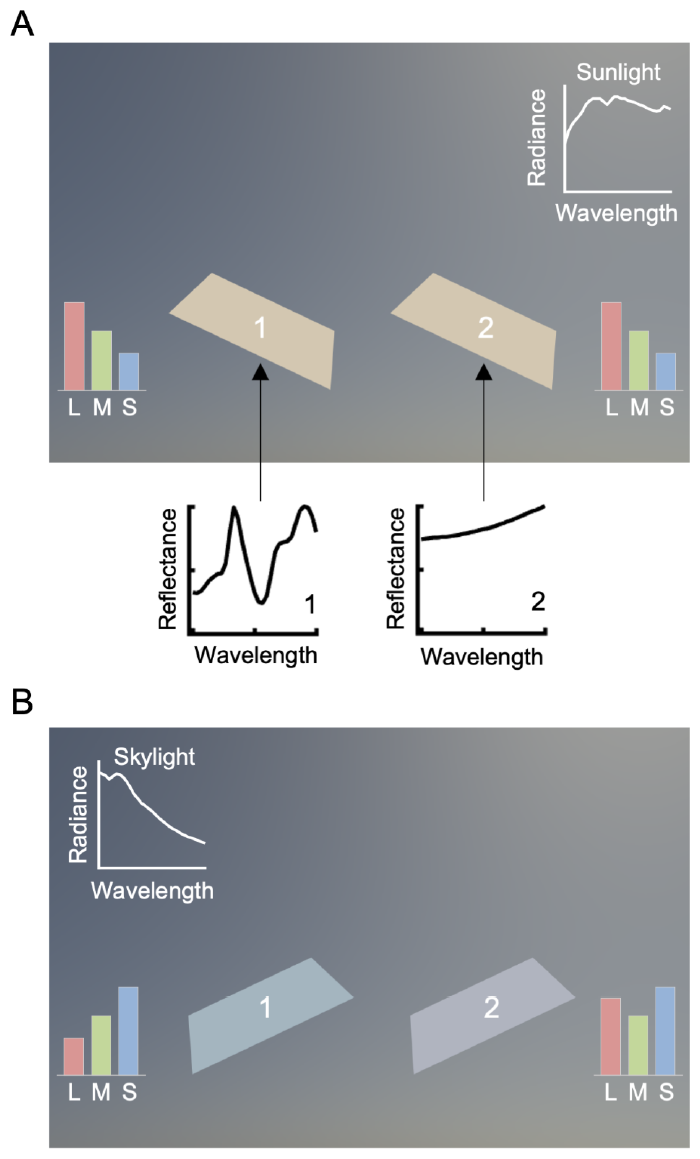
Two surfaces placed under directional lighting environments containing sunlight and skylight are tilted to mainly sample sunlight from the upper right corner in the environment (panel A). Their surface colors appear the same to human eyes even though they have distinct surface reflectances. However, by tilting the surfaces to sample skylight, these two surfaces become distinguishable (panel B).

## Results

Trichromats can solve 88.5% of metameric pairs in outdoor environments and 81.5% in indoor environments simply by tilting the surface (Figure 2, A and B). Dichromats can solve an even higher proportion (deutan: 93.6%, protan: 93.7%, tritan: 97.1% in outdoor scenes, and deutan: 85.8%, protan: 84.4%, tritan: 90.6% in indoor scenes). The main effects of scene type (outdoor or indoor) and colour vision type (trichromat, deutan, protan or tritan) were significant (*F*(1,8) = 5.40, *η*^*2*^ = 0.40, *p* = 0.0486; *F*(3, 24) = 18.8, *η*^*2*^ = 1.54, *p* < 0.00001) while the interaction between them was not (*F*(3,24) = 0.530, *η*^*2*^ = 0.0620, *p* = 0.666). Multiple comparisons using Bonferroni’s correction (significance level 0.05) found that the proportion was larger for any type of dichromat than for trichromats, and for tritan compared to deutan and protan. Thus, in real-world lighting environments, illuminant metamerism rarely causes issues for human observers.

**Figure 2:**
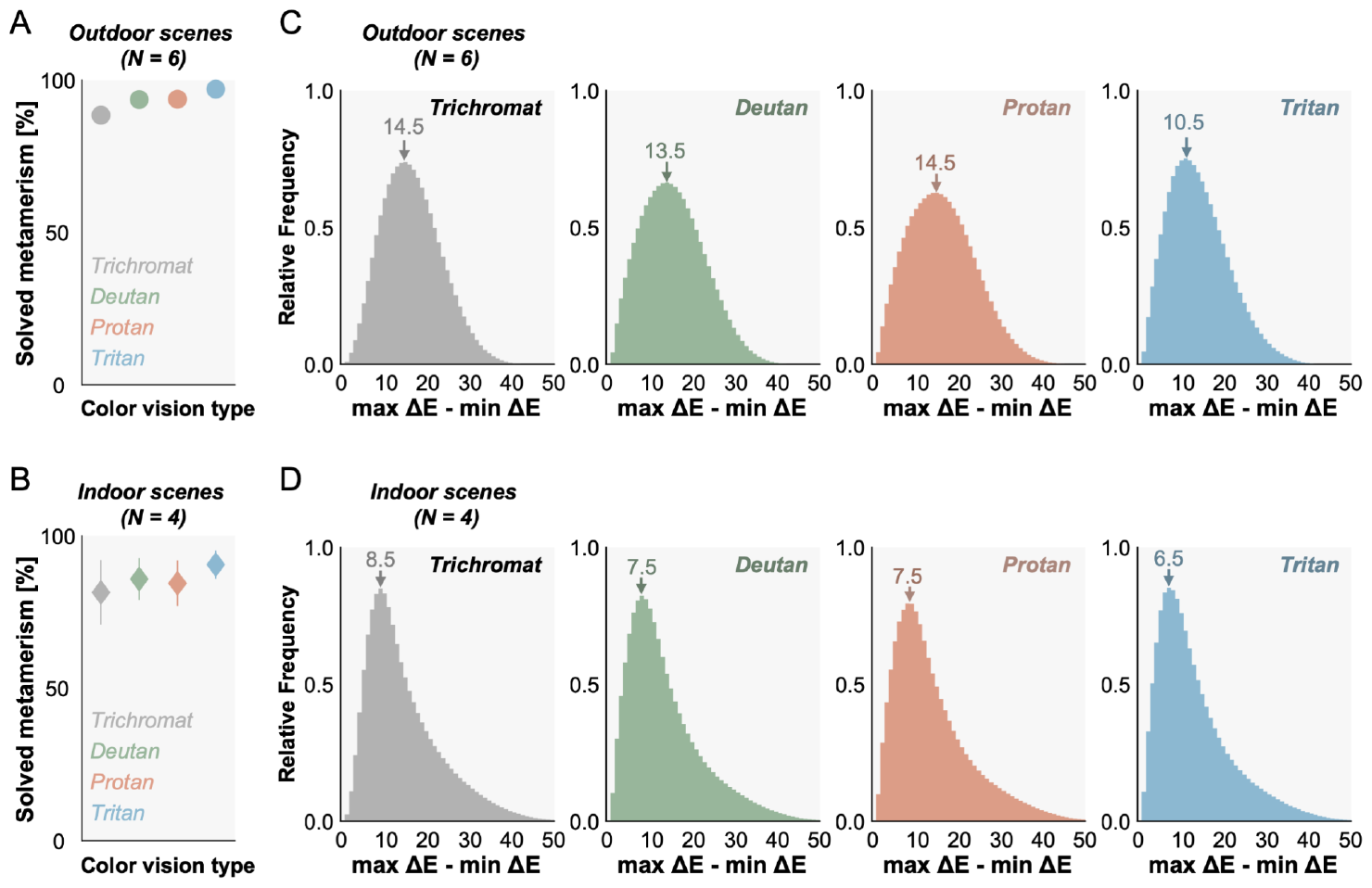
(A and B) The proportion of metamerism that can be solved by tilting a surface. The color depicts different color vision types. The values were averaged across 6 outdoor environments (A) and 4 indoor environments (B). Error bars show ± 1.0 S.D. across scenes. (C and D) Histograms of the absolute increase of ΔE defined by the difference between max ΔE and min ΔE, for outdoor scenes and indoor scenes, respectively. Data were pooled over 6 outdoor scenes or over 4 indoor scenes.

Here we used a criterion of ΔE = 0.36 in CIECAM02-UCS (Carter et al. 2018), which corresponds to the average radius of MacAdam ellipses in the color space (Martínez-García et al., 2013), as a criterion for ‘distinguishable’. Yet no official recommendation has been provided by the International Commission on Illumination (CIE). We thus checked the extent to which this finding holds across different criteria (ΔE = 0.5, 1.0 and 2.0). We additionally computed the proportion of broken metameric pairs at ΔE = 0.5, 1.0, and 2.0 for outdoor scenes and for indoor scenes. Overall the results suggest that the percentages slightly decrease as the ΔE values increase. By increasing the criterion ΔE from 0.36 to 2.0, the proportions of solved metamerism decrease from 88.5% to 82.1% for trichromats, 93.6% to 80.9% for deutan, 93.7% to 79.6% for protan and from 97.1% to 83.9% for tritan in outdoor scenes. Similarly for indoor scenes, the proportion decreases from 81.5% to 75.6% for trichromat, 85.8% to 73.0% for deutan, 84.4% to 72.4% for protan, and from 90.6% to 73.6% for tritan. However, importantly the general conclusion that a large proportion of metamerism can be solved still holds.

Sampling directional spectral variation in real environments yields a more general benefit since it always increases discriminability, regardless of the criterion for distinguishability. Thus, we also evaluated how much ΔE between pairs of surfaces increases by tilting the pairs. The mode increase of ΔE ranges between 10.5 and 14.5 for outdoor scenes and between 6.5 and 8.5 for indoor scenes (Figure 2, C and D), confirming that the discriminability of a pair of surfaces dramatically increases both for trichromats and dichromats.

These are the increases of ΔE in absolute terms, but the same absolute increase could convey different meanings. For example, an increase from 0.1 to 10.1 and an increase from 10 to 20 represent the same absolute improvement (i.e. plus 10) but relatively speaking the former case shows a more significant change. Thus, we also evaluated the increase in a proportional sense by computing (max ΔE - min ΔE) / min ΔE × 100. We again drew histograms of the metric for each lighting environment and took the mode value, and then computed average mode values across 6 outdoor scenes and 4 indoor scenes.

For outdoor scenes, the improvement was 63.5% for trichromats and higher for dichromats (deutan: 99.5%, protan: 85.5%, and tritan: 98.3%). Similarly for indoor scenes, trichromats showed 69.5% improvement and values for dichromats were higher than trichromats (deutan: 89.4%, protan: 86.1%, and tritan: 96.1%). The main effects was not significant for scene type (outdoor or indoor, F(1,8) = 0.01, *η*^*2*^ = 0.0007, *p* = 0.923) but significant for colour vision type (*F*(1,8) = 0.01, *η*^*2*^ = 0.0007, p < 0.00001). The interaction between them was not significant (*F*(3,24) = 0.91, *η*^*2*^ = 0.338, *p* = 0.451). Multiple comparisons using Bonferroni’s correction (significance level 0.05) found that the proportion was larger for any type of dichromat than for trichromats but there was no significant difference among the types of dichromats.

## Discussion

Illuminant metamerism arises from univariant cone responses. In this sense, it has been taken as a hard limit on human color vision. However, we show that directional spectral variation in natural environments allows us to distinguish most of the reflectance pairs that are indiscriminable under a single-illuminant. These results add a new perspective to classic trichromatic theory (Young, 1802) that serves as a basis of modern colour vision research. The term trichromacy may imply that we have access to only three sensory inputs to create color, but this is true only when we consider a snapshot of our external world. Instead, in daily lives cone signals associated with a particular surface change in a dynamic way. Even if we are missing one type of cone, we might recover the ‘missing’ dimension by using signals sampled at different time points (Broackes, 2010).

Precise measurements of directional spectral variation of the illumination in natural environments allowed us to perform the present analysis for the first time. We used an ideal model surface that does not capture all the complexity of natural objects. Moreover, the present study did not incorporate the complex behavior of human observers interacting with objects when discriminating surface colors. For example, dichromats might be motivated to explore surfaces in a different way from trichromatic observers, to improve their discrimination. Even considering these limitations, however, it is clear that the amount of extra information obtained just by tilting the surfaces is striking.

Some recent studies have suggested that illuminant metamerism is relatively infrequent when natural reflectances and illuminants are considered (Foster et al. 2006; Akbarinia & Gegenfurtner, 2018). Here we find that metamerism is even less frequent because of directional spectral variation in real world lighting. Quantification of such effects was made possible using new light-measurement data and computer-graphics rendering techniques that allow precise physical simulation of light-material interactions. It is striking that the complexity inherent in natural lighting environments can compensate for the biological limitations of trichromatic and dichromatic visual systems to a large extent.

## Materials and Methods

### Simulation using computer graphics techniques

We used computer-graphics techniques to evaluate the frequency of metamerism for planar surfaces placed in a three-dimensional environment defined by the measured illumination maps (Figure 3 left). The surface was matte (no specular reflection) and tiltable so that it could sample spectra from different directions in the scene. We simulated 20,132 surface reflectances measured from natural and manmade objects (see Supplementary information for details) and rendered the surface hyperspectrally (Jakob, 2010) at 81 tilts (Figure 3 right, grid images) under 10 different environmental illuminations. Depending on the tilt of the surface, the spectrum and chromaticity of the reflected light changes (Figure 3, rightmost plots). The resultant hyperspectral images were converted to XYZ 1931 images, then to CIECAM02-UCS images. We used Brettel’s model to convert XYZ tristimulus values for trichromats to XYZ for dichromats (Brettel et al., 1997). For each of 2.03×10^8^ reflectance pairs (20,132×20,131×0.5), we computed the chromatic difference ΔE at 81 tilts. We then counted the number of surface pairs that were indistinguishable (metameric) at least one tilt and calculated the proportion of these that became distinguishable at at least one other tilt (Figure 2 A, B). As a criterion of being distinguishable, we chose ΔE = 0.36 (Martínez-García et al., 2013). For each of the 2.03 × 10^8^ surface pairs we considered, we calculated the absolute increase of ΔE across 81 slants defined by computing maximum ΔE minus minimum ΔE. We performed this computation for all lighting environments and concatenated the data separately for outdoor scenes and for indoor scenes as shown in panels C and D in Figure 2. Thus, each histogram in panel C and panel D consists of 1.22 × 10^9^ (2.03 × 10^8^ × 6) and 8.12 × 10^8^ (2.03 × 10^8^ × 4) data points, respectively. We will provide a link to download hyperspectral environmental illuminations used for the simulations.

**Figure 3:**
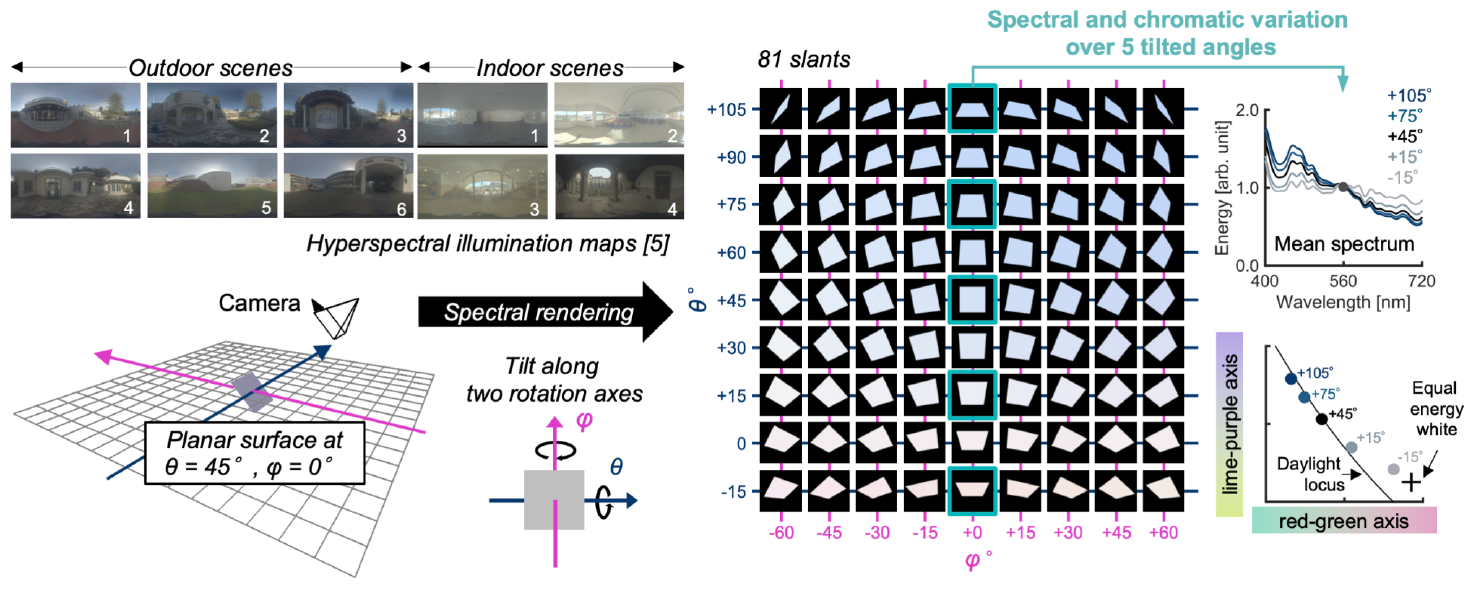
A three-dimensional scene in which a tiltable planar surface was placed. One of ten hyperspectral environmental illumination maps was applied to render the scene. For each of 20,132 surface reflectances, we hyperspectrally rendered the image of the tiltable surface at 81 tilts. The right-hand plots show that the mean spectrum and mean chromaticity change dramatically over the 5 selected tilts.

### Reflectance dataset

We first collected 54,282 reflectances measured from a variety of items: 2,471 flowers, 1,254 fruits, 8,051 human skin samples, 1,498 leaves, and 41,008 man-made objects (e.g. ink or dyes). They were obtained via publicly available datasets (Regan et al. 2018; Arnold et al. 2008; Matsumoto et al. 2014), purchased (Tajima et al., 1999) or via personal communication with Dr. Alexander Shenkin. However, there were many reflectance pairs that were physically very similar. This is problematic because in an extreme case, a pair of two identical surface reflectances is indistinguishable under any illumination, and thus using these spectra leads to an overestimation of the number of metameric pairs. To address this, we reduced the number of reflectances by identifying reflectance pairs that have a Pearson’s correlation coefficient of more than 0.999 in the spectral domain, and removing one reflectance from each such pair. After this procedure 20,132 reflectances (2,143 flowers, 991 fruits, 141 human skin, 1,164 leaves, and 15,693 man-made objects) remained, which were used for the analyses in this study.

### Perceptually uniform color space

Past studies have used a variety of metrics to estimate the frequency of metamerism. For example, one early study used Euclidian distance in XYZ tristimulus values (Stiles & Wyszecki, 1962), while a more recent study (Akbarinia & Gegenfurtner, 2018) chose to evaluate differences in a uniform colour space that accounts for the sensitivity of our visual system to colour differences in different regions of the space. We chose to evaluate differences between pairs of colored surfaces in a perceptually uniform color space called CIECAM02-UCS (Carter et al., 2018). There are several parameters that need to be defined to construct this colour space. Parameters were set separately for each lighting environment. For the white point Wp, we took the average XYZ1931 values across 81 rendered tilts of a surface that had a uniform reflectance (R=1.0 for all wavelengths). For the adapting field luminance LA, we used the mean XYZ1931 values across all pixels in the environmental illumination. For the relative luminance Yb, we used the ratio between the luminance of the adapting field (LA) and the luminance of the white point (Wp). These parameters were designed to incorporate the scene’s white point Wp, the luminance level of the background LA and observers’ adaptation state Yb that are known to modulate colour appearance. We set the viewing condition to “average” because this is often used as the condition appropriate for viewing the colour of a surface.

## Acknowledgments

The authors acknowledge Dr. Arash Akbarinia and Dr. Alexander Shenkin for helping us collect reflectance datasets. This work was supported by the Portuguese Foundation for Science and Technology (FCT) in the framework of the Strategic Funding UIDB/04650/2020. TM is supported by a Sir Henry Wellcome Postdoctoral Fellowship from Wellcome Trust (218657/Z/19/Z) and a Junior Research Fellowship from Pembroke College, University of Oxford. For the purpose of open access, the author has applied a CC BY public copyright license to any Author Accepted Manuscript version arising from this submission.

## Data availability

We will provide a link to download hyperspectral environmental illuminations used for the simulation at the time of publication.

